# Microbiota Accessible Carbohydrates Suppress *Clostridium difficile* in a Murine Model

**DOI:** 10.1101/243253

**Authors:** Andrew J Hryckowian, William Van Treuren, Samuel A Smits, Nicole M Davis, Jackson O Gardner, Donna M Bouley, Justin L Sonnenburg

**Author notes:** Correspondence Please address all correspondence to Justin Sonnenburg.

## Abstract

*Clostridium difficile* (*Cd*) is an opportunistic diarrheal pathogen and *Cd* infection (CDI) represents a major healthcare concern, causing an estimated 15,000 deaths per year in the United States alone^1^. Several enteric pathogens, including *Cd*, leverage inflammation and the accompanying microbial dysbiosis to thrive in the distal gut^2^. Although diet is among the most powerful available tools for affecting the health of humans and their relationship with their microbiota, investigation into the effects of diet on CDI has been limited. Here, we show in mice that the consumption of microbiota accessible carbohydrates (MACs) found in dietary plant polysaccharides has a significant impact on CDI. Specifically, using a model of antibiotic-induced CDI that typically resolves within 12 days of infection, we demonstrate that MAC-deficient diets perpetuate CDI. We show that *Cd* burdens are suppressed through the addition of either a diet containing a complex mixture of MACs or a simplified diet containing inulin as the sole MAC source. We show that switches between these dietary conditions are coincident with changes to microbiota membership, its metabolic output and *Cd*-mediated inflammation. Together, our data demonstrate the outgrowth of MAC-utilizing taxa and the associated end products of MAC metabolism, namely the short chain fatty acids (SCFAs) acetate, propionate, and butyrate, are associated with decreased *Cd* fitness despite increased *Cd* toxin expression in the gut. Our findings, when placed into the context of the known fiber deficiencies of a human Western diet, provide rationale for pursuing MAC-centric dietary strategies as an alternate line of investigation for mitigating CDI.

CDI is typically associated with antibiotic-mediated dysbiosis, yet 22% of individuals with community acquired CDI have no recent history of antibiotic use^3^. We and others previously demonstrated that direct microbiota-Cd metabolic interactions are critical determinants of *Cd* fitness in the distal gut^4-6^ and that the absence of dietary MACs leads to the expression of inflammatory markers by the host colonic epithelium^7^. Additional *in vitro* work suggested that MAC-centric metabolic interactions may play a role in reducing the fitness of *Cd* in the gut^8-10^, leading us to hypothesize that a MAC-deficient diet reinforces the inflammation and dysbiosis conducive to CDI.

We used an experimental model of CDI in ex-germ-free Swiss-Webster mice colonized with the microbiota of a healthy human donor (see Methods). These humanized mice were fed a diet containing a complex mixture of MACs (MAC_+_), or two diets that are both MAC-deficient (MD) (Fig. 1a). Mice fed the MD diets show persistent CDI while mice fed the MAC_+_ diet clear the pathogen below detection within 10 days of infection. After 36 days of persistent infection in mice fed the MD diets, a dietary shift to the MAC_+_ diet results in clearance below detection within 9 days (Fig. 1a). This MAC_+_ diet-mediated CDI suppression is also observed in conventional C57BL/6 and Swiss-Webster mice and in ex-germ-free Swiss-Webster mice colonized with a conventional Swiss-Webster mouse microbiota (Fig. S1), demonstrating that MAC-dependent CDI suppression is not confined to a specific microbiota or host genotype.

**Figure 1.**
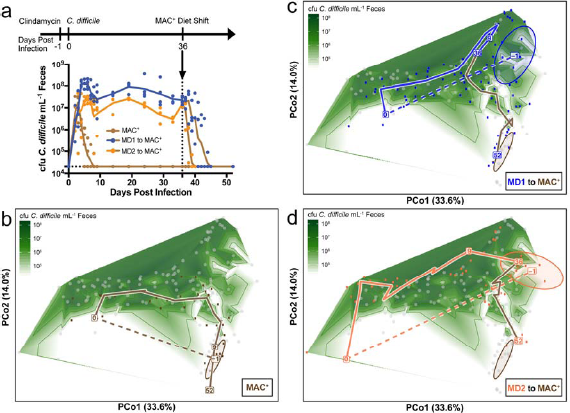
Dietary MACs toggle the fitness of *Clostridium difficile (Cd)* in the gut while engendering distinct microbiota states. Humanized, age matched female Swiss-Webster mice were maintained on a diet containing a complex mixture of MACs (MAC^+^, n=4 mice) or on diets deficient in MACs (MD1, n=4 mice; and MD2, n=4 mice) starting 8 days pre-infection. See Methods for details of diet compositions. (a) Mice were subsequently gavaged with clindamycin at 1 day before infection with *Cd.* At 36 days post-infection, mice fed the MD diets were switched to the MAC^+^ diet. *Cd* burdens were monitored over time by selective culture, as described in Methods. One of the MD2 fed mice was moribund on D10 and was euthanized. Individual per-sample *Cd* burdens are plotted and lines represent geometric mean burdens per time point. (**b-d**) Principal coordinates analysis plots of Weighted UniFrac distances between microbiota samples collected from these mice were prepared and overlaid with log-fold contour plots of *Cd* burdens, as measured in panel a. Clindamycin affects the composition of the microbiota in all groups of mice from D-1 (annotated with ellipses representing 80% CI) to D0 (dotted lines), resulting in a dysbiotic state permissive to CDI. In each panel, a line is drawn through the centroid of the points for a given experiment day. (c) In mice fed the MAC_+_ diet, the microbiota returns to resemble the pre-infection state as *Cd* burdens decrease. (d, e) In mice fed the MD1 and MD2 diets, respectively, CDI remains unresolved until dietary intervention with the MAC_+_ diet at D36, which shifts the microbiota to resemble that of other MAC^+^ fed mice as *Cd* burdens decrease (brown lines). Points are colored by the highlighted treatment group, or alternatively are retained as gray points for reference. See **Fig. S4** for further explanation on generation and interpretation of the contoured PCoA (cPCoA) plots shown in panels b-d.

To enumerate gut-resident microbes that might suppress *Cd,* we sequenced 16S rRNA amplicons from the feces of humanized mice (Fig. 1a). The presence of dietary MACs and treatment with antibiotics affected both alpha and beta diversity of operational taxonomic units (OTUs) in the gut microbiota (Figs. 1b-d, S2, S3). Two principle coordinates explain 48% of the variance in weighted UniFrac distances between samples. We traced temporal changes in the composition of the microbiota through this space. To highlight the composition of the microbiota in the context of CDI, a log-fold contour plot was drawn to illustrate burdens of *Cd* that correspond to these samples (see Fig. S4 for further explanation of these “contoured PCoA” [cPCoA] plots).

Clindamycin remodels the mouse microbiome to a Cd-permissive state (Figs. 1b-d, dotted lines; Fig. S3). After inoculation with *Cd,* the microbiota of MAC_+_-fed mice changes significantly, and as *Cd* burdens decrease, the community returns to resemble the pre-infection state (Figs. 1b, S3a), illustrating compositional resilience of the microbiota under the MAC_+_ dietary condition during CDI. During persistent infection in mice fed the MD diets, the microbiota composition is similar to the pre-infection MAC-deficient-associated microbiota. However, after the MAC_+_ dietary intervention, the microbiota of these mice transitions to resemble the microbiota of mice prophylactically fed the MAC_+_ diet as *Cd* burdens decrease (Figs. 1c, 1d, S3b, S3c). These data suggest that diet and antibiotic treatment are two major drivers of microbial communities that support or exclude *Cd* in our model. Furthermore, the similarities in the microbiota of uninfected and persistently infected mice fed the MD diets may be due to the metabolic and compositional constraints imposed by this dietary condition, which we hypothesize is supportive to *Cd* during infection.

Having shown that the MAC_+_ diet, containing a complex and ill-defined mixture of MACs (see Methods), is successful in suppressing CDI, we sought to decouple the effects of MACs from other dietary components (e.g. phytonutrient^11^ or protein^12^ content). Using a simplified MD1-based diet containing inulin as the sole MAC source, the findings from our initial dietary intervention experiment in Cd-infected animals are recapitulated (Figs. 2a, S5, S6, Supplemental Text 1). Furthermore, prophylactic inulin feeding (either 10% in the diet as above or 1% in the drinking water of mice fed the MD1 diet), results in dose-dependent effects on both the maximum *Cd* burden and on *Cd* clearance kinetics (Fig. S7). Taken together, like the complex MAC_+_ diet, inulin feeding reduces *Cd* burdens across experimental paradigms (Figs. 2a, S6-S8, Supplemental Text 2).

**Figure 2.**
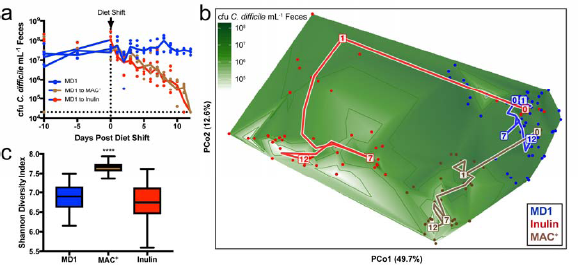
A diet containing inulin as the sole MAC source reduces Cd burdens without increasing microbial diversity. Mice with persistent CDI were subjected to a dietary intervention of the complex MAC^+^ diet (n=3), a diet containing inulin as the sole MAC source (n=4), or were maintained on the MD1 diet (n=4). See Methods for details of diet compositions. (a) Burdens of *Cd* were monitored over time by selective culture as described in Methods. Individual per-sample *Cd* burdens are plotted and lines represent geometric mean burdens per time point. (b) A contoured PCoA (cPCoA) plot of weighted UniFrac distances between microbiomes of these mice was prepared and overlaid with log-fold contour plots of *Cd* burdens, as measured in panel A. A line is drawn through the centroid of the points for a given experimental day. See **Fig. S4** for further explanation on generation and interpretation of cPCoA plots. (c) Alpha diversity of communities was determined longitudinally for the microbiota of these mice by Shannon Diversity Index. Differences in *Cd* burdens and alpha diversity between dietary conditions was determined by collapsing all time points after the diet shift time point (See **Fig. S11** for temporal differences in Shannon Diversity Index). Statistical significance was determined by Kruskal Wallis test with Dunn’s multiple comparison test (****=p<0.0001).

Although the complex MAC_+_ and the inulin-containing diets both negatively impact the *in vivo* fitness of *Cd,* the overall microbiome composition differs between mice fed these two diets, as illustrated by a two-dimensional cPCoA subspace that explains 62% of the variation in the data and the proportional abundance of taxa (Figs. 2b, S9, S10). Because increased gut microbiota diversity is associated with resistance to a number of pathogens and is a hallmark of FMT-mediated suppression of CDI^6,13-15^, one hypothesis is that MACs positively affect CDI outcome by favoring a diverse microbiota. Though CDI suppression induced by the MAC_+_ diet is correlated with an increase in alpha diversity of the gut microbiota, alpha diversity does not increase when *Cd* burdens decrease via inulin feeding (Figs. 2c, S11a). Furthermore, community evenness is lowest in the inulin fed mice (Fig. S11b), consistent with a limited number of taxa profiting from a single MAC type^16^. Together, these data demonstrate that MACs significantly reduce *Cd* burdens but that this reduction is independent of microbiota diversity.

Despite these wholesale differences in community composition, we pursued two of several possibilities: (1) a common subset of OTUs suppress CDI across dietary interventions or (2) diet-specific but functionally similar OTUs within dietary conditions suppress CDI (see Supplemental Text 3, Tables S1, S2). Notably, several taxa are significantly (anti)correlated with *Cd* burdens regardless of diet, suggesting that common microbial signatures underlie permissive and non-permissive states (Table S3). Among these, *Parabacteroides,* Lachnospiraceae, and Erysipelotrichaceae are correlated with *Cd* abundance regardless of diet. The correlation between *Parabacteroides* and *Cd* burden is consistent with previous observations that *Parabacteroides* are elevated in *Cd “*supershedder” mice^17^. In humans, Erysipelotrichaceae and some Lachnospiraceae are enriched in individuals with CDI compared to nondiarrheal controls^18^. Despite these commonalities, the majority of features identified in Table S3 are (anti)correlated with *Cd* in a subset of diets, supporting previous work that distinct context-dependent communities, rather than core Cd-(un)supportive communities, are important for determining CDI status^19^. Because others have demonstrated that metabolites, rather than microbes, are able to differentiate CDI status in humans^20^, we hypothesized that diet creates metabolic landscapes that are either supportive or unsupportive of *Cd*.

Therefore, we pursued whether MAC-mediated suppression of CDI could be differentiated from the permissive condition on a molecular basis relevant to MAC metabolism. We measured the major metabolic end products of MAC metabolism (the SCFAs acetate, propionate, and butyrate) in cecal contents of mice (see Fig S6). Acetate and butyrate are elevated in the ceca of mice fed MAC_+_ and inulin diets relative to those fed the MD1 diet, and propionate is elevated in the ceca of MAC_+_-fed mice relative to those fed the MD1 or inulin diets (Fig. 3a). Furthermore, acetate, propionate, and butyrate have concentration-dependent negative effects on *Cd* growth as measured by differences in doubling time (Fig. 3b) and concentration-dependent positive effects on expression of a critical *Cd* virulence factor, the glycosylating toxin TcdB^21^ (Fig. 3c). Our findings using *Cd* strain 630 are consistent with previous findings in other *Cd* strains: *(i)* SCFAs inhibit the growth of five non-630 strains in a concentration dependent fashion^10,22^ and *(ii)* butyrate affects toxin expression in *Cd* strain VPI 10463^23^.

**Figure 3.**
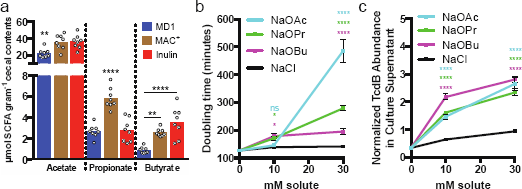
Acetate, propionate, and butyrate are elevated in the ceca of mice fed MACs, and affect growth and toxin production in *C. difficile.* (a) The short chain fatty acids acetate, propionate, and butyrate were measured in cecal contents of mice fed the MD1, MAC^+^, and inulin containing diets (n=8 per dietary condition). See **Fig S6** for time points of sacrifice corresponding to measurements). Data points represent per sample concentration of each SCFA and bars represent the mean per-diet SCFA concentration. Statistical significance was assessed by ordinary one-way ANOVA and Tukey’s multiple comparison test. (**, p<0.01; ****, p<0.0001) (b) Doubling time was calculated for *Cd* grown in CDMM supplemented with 0, 10, and 30 mM sodium acetate (NaOAc), sodium propionate (NaOPr), sodium butyrate (NaOBu), or sodium chloride (NaCl, sodium matched controls) as described in Methods. For each growth condition, at least n=16 independent cultures were assessed. (c) TcdB was quantified in supernatants from n=12 independent cultures from each growth condition as described in Methods. Points represent mean values, error bars represent ±SEM (panels b-c). Statistical significance was assessed for each SCFA and sodium matched controls at each concentration via Mann-Whitney test (ns, not significant; *, p<0.05; ****, p<0.0001).

Given these findings, we hypothesize that dietary MACs negatively affect the fitness of *Cd* in two interrelated ways. First, MACs drive privileged outgrowth of MAC-utilizing members of the microbiota (e.g. *Bacteroides* spp., see Table S2b). Second, the SCFAs that result from MAC metabolism negatively affect the fitness of *Cd,* which could be due to the buildup of endproducts of key metabolic pathways, such as reductive acetogenesis and butyrogenesis^5,24^. The expression of TcdB (and the co-regulated toxin TcdA) in *Cd* is controlled by multiple inputs, such as nutrient availability, quorum sensing, and other environmental stresses^25^. Therefore, it is possible that SCFAs serve as a signal to *Cd* of microbiome fermentation, and this signal of a competitive and inhospitable gut environment leads to an increase in toxin production.

Since limitation of dietary MACs is known to increase inflammation in the gastrointestinal tract^7^, we examined colonic histopathology and toxin levels to better understand the inflammation during MAC-dependent suppression of CDI. Humanized Swiss-Webster mice fed the MD1 diet were infected as in Fig. 1a and were switched to either the MAC_+_ or inulin diet at 7 days post infection. At pre- and post-diet shift time points, histopathology of proximal colon tissue was evaluated. Pathology was significantly increased in all infected mice relative to uninfected control mice fed the MAC_+_ or inulin diets (Figs. 4a, S12; Table S4). Notably, inflammation is comparably elevated in both infected and uninfected mice fed the MD1 diet, consistent with the contribution of the MD diets to inflammation and *Cd* persistence.

**Figure 4.**
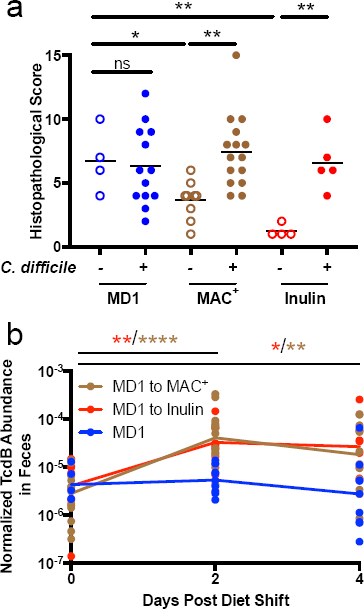
Inflammation and *Cd* toxin expression are diet-dependent. Age-matched, female, humanized Swiss-Webster mice were fed the MD1 diet, gavaged with clindamycin, and subsequently infected with *Cd* as in Fig. 1 or were mock infected with filter sterilized PBS. After 7 days of infection, mice were switched either to the MAC^+^ or inulin-containing diet. (a) Mice were euthanized before and after diet change, at time points specified in Table S4. Histopathology was carried out on proximal colon tissue from these mice as described in Methods (n=4 for mock-infected mice fed the MD1 diet, n=9 for mock-infected mice fed the MAC^+^ diet, n=4 for mock-infected mice fed the inulin diet, n=13 for infected mice fed the MD1 diet, n=15 for infected mice fed the MAC^+^ diet, and n=5 for infected mice fed the inulin diet). Individual points represent the sum histopathological score and lines are drawn at the mean score for each group. Statistical significance between relevant groups was assessed by unpaired T test (*= p<0.05; **=p<0.01). (b) For infected mice where >5 mg fecal material could be collected (n=14, n=13, and n=13 for mice at 0, 2, and 4 days post MD1 to MAC^+^ diet shift; n=8, n=9, and n=6 for for mice at 0, 2, and 4 days post MD1 to inulin diet shift; and n=5, n=9, and n=9 at matched time points for mice that were maintained on the MD1 diet), levels of TcdB in the feces were measured and normalized to the burdens of *Cd* detected. Points represent normalized TcdB abundance for individual fecal samples lines are drawn through the per-day per-diet geometric mean normalized toxin abundance. For mice switched to the MAC^+^ diet, normalized TcdB abundance increases 14.5-fold from day 0 to day 2 (p<0.0001) and 6.5-fold from day 0 to day 4 (p=0.0066). For mice switched to the inulin diet, normalized TcdB abundance increases 8.0-fold from day 0 to day 2 (p=0.0016) and 6.5-fold from day 0 to day 4 (p=0.0426). Statistical significance between relevant pairs of treatment groups was assessed by Mann-Whitney test (*= p<0.05; **=p<0.01; ***= p<0.001; ****=p<0.0001).

We also measured burdens of *Cd* and levels of TcdB in feces during the shift from the MD1 diet to either the MAC_+_ or inulin diets. TcdB is detected during persistent infection but its expression is further elevated at 2 and 4 days after the shift to the MAC_+_ or inulin diet (Fig. 4b). Though cfu-normalized TcdB abundance is elevated after the diet shift, the overall abundance of TcdB decreases from day 2 to day 4 post diet shift (Fig. S13). This shows that toxin expression is elevated on a per-cell basis in response to MACs but that overall TcdB abundance scales with decreasing burdens of *Cd.* Together, these data support a model where MAC deficient diets facilitate a level of inflammation supportive of *Cd* survival in the gut, enabling *Cd* persistence despite lower levels of toxin expression. *Cd* responds to the non-permissive MAC-driven environmental change by elevating toxin expression. However, *Cd* is unable to maintain or regain its niche in the gastrointestinal tract upon the sustained consumption of MACs by the host.

In the context of CDI, the relative contribution of inflammation to the exclusion of inflammation-sensitive competitors versus the creation of privileged nutrients, analogous to strategies delineated for the enteric pathogen *Salmonella* Typhimurium, remains to be determined^2^. Furthermore, though SCFAs can directly inhibit bacterial growth and affect *Cd* toxin production, additional work is needed to investigate the potential roles of other relevant factors that could be impacting infection dynamics. For example, SCFA can signal through host pathways (e.g. improved barrier function via hypoxia-inducible factor^26^) which may influence inflammation and microbiome composition independently of *Cd* burdens and toxin production; other metabolites such as bile acids can influence CDI^6^ and may interact with dietary effects.

Current microbiota-centric therapies for CDI, such as fecal microbiota transplant and probiotic administration, focus on the introduction of exogenous organisms. Our work in mice shows that dietary intervention supports microbial communities that exclude *Cd* without the requirement for microbe introduction. Regardless of the inter-experiment and inter-animal variations in clearance kinetics (Fig. S8), the effect is highly reproducible. Clearance kinetics may be further affected by host genetics, initial microbiota composition, or overall dietary MAC concentration/composition.

Despite the individuality in the gut microbiota of patients with CDI, there is a consistent metabolic response that underlies CDI across individuals^20^. In light of observations that MACs profoundly alter the composition and function of the microbiota and host physiology^27^, our findings raise the possibility that SCFAs are a critical part of the metabolic landscapes tied to CDI status across individuals. Currently, several technical hurdles exist to experimentally manipulating SCFAs within the colon, presenting a tremendous opportunity to develop new methods that enable exploring this important class of fermentation end-products and their effects on gut ecology (**Supplemental Text 4**).

Notably, two independent human trials have shown cooked green bananas (rich in MACs as evident by elevated SCFAs in the stool of treated patients) aid host recovery from another enteric pathogen, *Shigella*^28,29^. More recently, it was shown that a MAC-deficient diet leads to microbiota-dependent mucus degradation and attachment-dependent lethal colitis by the murine pathogen, *Citrobacter rodentium^30^.* Taken together, our work is part of a growing body of literature providing evidence that dietary manipulation of the metabolic networks of the intestinal tract is a powerful and underexplored way to influence gastrointestinal pathogens.

## Methods

### Media and bacterial growth conditions

Frozen stocks of *Clostridium difficile (Cd)* strain 630^31^ were maintained under anaerobic conditions in septum-topped vials. *Cd* 630 was routinely cultured on CDMN agar, composed of *Cd* agar base (Oxoid) supplemented with 7% defibrinated horse blood (Lampire Biological Laboratories), 32 mg/L moxalactam (Santa Cruz Biotechnology), and 12 mg/L norfloxacin (Sigma-Aldrich) in an anaerobic chamber at 37° (Coy). After 16-24 hours of growth, a single colony was picked into 5 mL of pre-reduced Reinforced Clostridial medium (RCM, Oxoid) and grown for 16 hours. This 16-hour culture was used to inoculate mice, below.

For in vitro experiments, *Cd* 630 was cultured on CDMN as above. Single colonies were picked into pre-reduced *Cd* minimal medium (CDMM) without glucose, as described previously^32^. After 16 hours of growth, subcultures were prepared at a 1:200 dilution in pre-reduced CDMM supplemented with 0, 10, or 30 mM of sodium acetate (Fisher), sodium propionate (Sigma Aldrich), sodium butyrate (Sigma Aldrich), or sodium chloride (EMD Millipore) in sterile polystyrene 96 well tissue culture plates with low evaporation lids (Falcon). To further minimize evaporation of culture media during growth, the 36 wells along the perimeter of the 96 well plates were filled with water rather than culture. Cultures were grown anaerobically as above in a BioTek Powerwave plate reader. At 15-minute intervals, the plate was shaken on the ‘slow’ setting for 1 minute and the optical density (OD_600_) of the cultures was recorded using Gen5 software (version 1.11.5). After 24 hours of growth, culture supernatants were collected after centrifugation (5 minutes at 2,500 x g) and stored at −20°C for quantification of TcdB (see **Quantification of *C. difficile* toxin TcdB**, below).

### Murine model CDI

All animal studies were conducted in strict accordance with the Stanford University Institutional Animal Care and Use Committee (IACUC) guidelines. Murine model CDI was performed on age- and sex-matched mice between 8 and 17 weeks of age, possessing one of three gut microbiota colonization states: (1) Humanized Swiss-Webster mice: Germ free mice (SWGF, Taconic; bred in house) were inoculated with a fecal sample obtained from a healthy anonymous donor, as used in previous studies from our laboratory^33,34^, (2) Conventionally-reared Swiss-Webster mice (SWRF, Taconic; bred in house); or C57BL/6 mice (B6EF, Taconic; experiments conducted on animals acquired directly from vendor), and (3) Conventionalized mice: Germ free Swiss-Webster mice were inoculated with a fecal sample obtained from SWRF mice. The gut microbiomes of the humanized and conventionalized mice were allowed to engraft for at least 4 weeks^35^.

To initiate CDI, mice were given a single dose of clindamycin by oral gavage (1 mg; 200 μL of a 5 mg/mL solution) and were infected 24 hours later with 200 μL of overnight culture grown in RCM (approximately 1.5×10^7^ cfu/mL) or mock infected with 200 μL filter sterilized PBS. To reactivate CDI in mice that had cleared the infection below detection, mice were given a single dose of clindamycin as above.

Feces were collected from mice directly into microcentrifuge tubes and placed on ice. To monitor *Cd* burdens in feces, 1 μL of each fecal sample was resuspended in PBS to a final volume of 200 μL, 10-fold serial dilutions of fecal slurries (through 10^−3^-fold) were prepared in sterile polystyrene 96 well tissue culture plates (Falcon). For each sample, duplicate 10 μL aliquots of each dilution were spread onto CDMN agar. After 16-24 hours of anaerobic growth at 37°C, colonies were enumerated and duplicate spots were averaged to give cfu values (limit of detection = 2×10^4^ cfu/mL feces). *Cd* was undetectable in all mice prior to inoculation with *Cd* (Figs. 1A, S1, S6-S8) and in all mice that were mock infected with PBS (Fig. S8), supporting that the animals used in this work were not pre-colonized with *Cd* (e.g. *Cd* LEM1, as seen by Etienne-Mesmin and colleagues^36^). After serial dilution of fecal samples, the remaining amounts of fecal samples were immediately frozen at −80°C until needed for 16S rRNA analysis and TcdB ELISAs, below. It was not possible to blind researchers to infection or dietary status of the animals.

### Mouse diets

Mice were fed one of four diets in this study ad libitum: (1) a diet containing a complex mixture of MACs (MAC_+_, Purina LabDiet 5010); (2) a custom MAC-deficient diet^37^ [MD1, 68% glucose (w/v), 18% protein (w/v), and 7% fat (w/v) (Bio-Serv); (3) a commercially available MAC-deficient diet [MD2, 34% sucrose (w/v), 17% protein (w/v), 21% fat (w/v); Harlan TD.88137], or (4) a custom diet containing inulin as the sole MAC source^37^, which is based on the MD1 diet [58% glucose (w/v); 10% inulin (w/v) [Beneo-Orafti group; OraftiHP]; 18% protein (w/v), and 7% fat (w/v) (Bio-Serv)]. Where applicable (see Fig. S7), the drinking water of mice fed the MD1 diet was supplemented with 1% inulin (w/v) [Beneo-Orafti group; OraftiHP]. Because mice consume approximately 5 grams of food per day and 5 mL of water per day^38^, water with 1% inulin gives an approximate 10-fold reduction in inulin consumed relative to the 10% inulin diet. Where applicable (see Fig. S14), mice fed the MD1 diet were gavaged daily with 200 μL tributyrin (Sigma Aldrich) or were given a cocktail of sodium acetate (67.5 mM, Fisher), sodium propionate (25 mM, Sigma Aldrich), and sodium butyrate (40 mM, Sigma Aldrich) in their drinking water^39^ starting 1 day pre-infection. Groups of mice were randomly assigned to dietary conditions.

### Histology and histopathological scoring

Proximal colon sections harvested for histopathologic analyses were fixed in 10% buffered neutral formalin and routinely processed for paraffin embedding, sectioned at 4 microns, mounted on glass slides and stained with hematoxylin and eosin (Histo-tec Laboratory, Hayward, CA). Analyses were performed by a board certified veterinary pathologist, using a semiquantitative scoring system^40^ that evaluated distribution and severity of cellular infiltrates (inflammation), gland hyperplasia, edema, and epithelial disruption using a severity score of 0 to 5 (0 = no significant lesion, 1 = minimal, 2 = mild, 3 = moderate, 4 = marked, 5 = severe, see Table S4).

### Quantification of *C. difficile* toxin TcdB

Levels of TcdB in culture supernatants and feces were quantified relative to a standard curve of purified TcdB using the “Separate detection of *C. difficile* toxins A and B” kit (TGC Biomics) according to the instructions that are readily available on the manufacturer’s website. For culture supernatants which were harvested at 24 hours post-inoculation (see Media and bacterial growth conditions, above), toxin abundance was normalized by the final OD_600_ of the culture and “Normalized TcdB Abundance in Culture Supernatant” = [(x ng toxin) / (y OD600)] and is reported in Fig. 3c. For each fecal sample, toxin abundance was normalized by the number colony forming units, as determined by selective culture on CDMN agar (above), for each sample; “Normalized TcdB Abundance in Feces” = [(x ng toxin gram^−1^ feces) / (y cfu *C. difficile* mL^−1^ feces)] and is reported in Fig. 4b. Non cfu-normalized TcdB abundance (ng toxin gram^−1^ feces) in mouse feces is reported in Fig. S13.

### 16S rRNA amplicon sequencing and OTU picking methods

Total DNA was extracted from frozen fecal material using the PowerSoil DNA Isolation Kit (MoBio) or the Powersoil-htp 96 well DNA isolation kit (MoBio). Barcoded primers were used to amplify the V3-V4 region of the 16S rRNA gene from extracted bacterial DNA using primers 515f and 806rB via PCR^41^. Amplicon cleanup was performed using the UltraClean PCR Clean-Up Kit (MoBio) and quantification was performed using the high sensitivity Quant-iT dsDNA Assay Kit (Thermo Fisher). Amplicons were pooled to an equimolar ratio. Amplicons from 3 mouse experiments were sequenced in 3 different paired-end Illumina MiSeq runs, with each experiment occurring on a separate run. The sample split/run corresponds to the field ‘Experiment’ in Table S1.

For commands executed for the 16S rRNA-based bioinformatics analysis, please see Code S1, an ipython notebook. Runs were demultiplexed independently due to some non-unique barcodes, and then concatenated prior to OTU picking using ‘split_libraries_fastq.py’ with default quality parameters in QIIME 1.9.1^42^. Open reference OTU picking was conducted with default parameters using the QIIME script ‘pick_open_reference_otus.py’ (with default clustering algorithm UCLUST^43^) on the 24,582,127 reads that passed quality filtering. OTUs whose representative sequence failed to align to the Greengenes reference alignment with at 85% identity using PyNAST were discarded^44,45^.

We removed OTUs occurring in at least 10 samples, and/or having less than 26 counts in the entire dataset. This filtering reduced the number of OTUs by 95.04% (211,884 to 10,504) but removed 5.2% of the feature-mass (23,293,178 to 22,078,743). This type of filtering removes a vast number of features that are likely artefacts, boosts power by reducing false discovery penalties, and concentrates analysis on biologically meaningful features. We rarefied our data to correct for differences sequencing depth. To ensure our results were not artefacts of rarefaction depth we conducted analyses at multiple rarefaction levels and our conclusions were not changed. We use OTU tables rarefied to 7,000 in this study, facilitating inter-run comparisons.

### Supervised learning

Using the ‘supervised_learning.py’ script from QIIME 1.9.1, the random forests classification method (with 10-fold cross validation error estimation) was trained using an OTU table as prepared above in “16S sequencing and OTU picking methods.” Presence or absence of *Cd* in a fecal sample (as determined by selective culture) or current diet were used as the class label category, corresponding to the field ‘Plus_minus_Cd’ and ‘Current_diet’ of Table S1. The OTU table was modified for this analysis by querying each of the 11 *Cd* 630 rRNA sequences against a BLAST database built from the representative set of OTUs created during OTU picking (see “16S sequencing and OTU picking methods”), after which the OTUs that matched *Cd* 630 rRNA sequences (cutoff 97% identity) were collapsed into a single *Cd* OTU,

“k_—_Bacteria;p_—_Frimicutes;c_—_Clostridia;o_—_Clostridiales;f_—_Peptostreptococcaceae;g_–_Clostridioides;s_—_putative_–_difficile.” See **Code S1** for the code used for this analysis.

### Quantification of short chain fatty acids (SCFAs)

Immediately following euthanasia at 32 days post infection, cecal contents were removed from mice described in **Fig. S6**, weighed, and flash frozen in liquid nitrogen. Cecal contents (70-150 mg) were suspended in a final volume of 600 μl in ice-cold ultra pure water and blended with a pellet pestle (Kimble Chase) on ice. The slurry was centrifuged at 2,350 x g for 30 seconds at 4°C and 250 μL of the supernatant was removed to a septum-topped glass vial and acidified with 20μL HPLC grade 37% HCl (Sigma Aldrich). Diethyl ether (500 µL) was added to the acidified cecal supernatant to extract SCFAs. Samples were then vortexed at 4°C for 20 minutes on ‘high’ and then were centrifuged at 1,000 x g for 3 minutes. The organic phase was removed into a fresh septum-topped vial and placed on ice. Then, a second extraction was performed with diethyl ether as above. The first and second extractions were combined for each sample and 250 µL of this combined solution was added to a 300 µL glass insert in a fresh glass septum-topped vial containing and the SCFAs were derivitized using 25 µL N-tert-butyldimethylsilyl-N-methyltrifluoroacetamide (MTBSTFA; Sigma Aldrich) at 60°C for 30 minutes.

Analyses were carried out using an Agilent 7890/5975 single quadrupole GC/MS. Using a 7683B autosampler, 1 µL split injections (1:100) were made onto a DB-5MSUI capillary column (30 m length, 0.25 mm ID, 0.25 µm film thickness; Agilent) using helium as the carrier gas (1 mL/minute, constant flow mode). Inlet temperature was 200°C and transfer line temperature was 300°C. GC temperature was held at 60°C for 2 minutes, ramped at 40°C/min to 160°C, then ramped at 80°/min to 320°C and held for 2 minutes; total run time was 8.5 minutes. The mass spectrometer used electron ionization (70eV) and scan range was m/z 50-400, with a 3.75-minute solvent delay. Acetate, propionate, and butyrate standards (20 mM, 2 mM, 0.2 mM, 0.02 mM, 0 mM) were acidified, extracted, and derivatized as above, were included in each run, and were used to generate standard curves to enable SCFA quantification.

### Measurement of doubling time for in vitro growth experiments

Raw OD_600_ measurements of cultures grown in CDMM (see “Media and bacterial growth conditions,” above) were exported from Gen5 to MATLAB and analyzed using the growth_curve_analysis_v2_SCFA.m script and analyze_growth_curve_SCFA.m function (Code S2 and Code S3, respectively). Growth rates were determined for each culture by calculating the derivative of natural log-transformed OD_600_ measurements over time. Growth rate values at each time point were then smoothed using a moving average over 75-minute intervals to minimize artefacts due to noise in OD measurement data. To mitigate any remaining issues with noise in growth rate values, all growth rate curves were also inspected manually. Specifically, in cases where the analyze_growth_curve_SCFA function selected an artefactual maximum growth rate, the largest local maximum that did not correspond to noise was manually assigned as the maximum growth rate. Doubling time was then computed by dividing the natural log of 2 by maximum growth rate. The investigator that conducted growth curve analysis was blinded to the experimental conditions in which growth curve data were obtained.

### Statistical methods

Alpha and beta diversity, correlations, and random forests were computed using QIIME (‘alpha_diversity_through_plots.py’, ‘beta_diversity_through_plots.py’, ‘observation_metadata_correlations.py’, ‘supervised_learning.py’). Kruskal-Wallis, Mann-Whitey, Student’s T, ANOVA, and D’Agostino-Pearson tests were performed using standard statistical analyses embedded in the Prism 7 software package (GraphPad Software Inc.). Spearman correlations were calculated in Python using **Code S1** under the heading ‘feature correlations by diet’. Specific statistical tests are noted in figure legends or tables as applicable.

## Data availability

The data that support the findings of this study are available from the corresponding author upon request. The 16S sequence data is uploaded to Qiita (http://qiita.ucsd.edu; Study ID 11347).

## Code availability

For custom code used in this study, see **Code S1-S3**.

## Acknowledgements

We thank Steven K. Higginbottom (Department of Microbiology and Immunology, Stanford) for expertise and technical assistance in all mouse experiments, Allis S. Chien (Stanford University Mass Spectrometry Facility) for developing the GC-MS parameters used in this study, and Carlos G. Gonzalez (Department of Chemical and Systems Biology, Stanford) for assistance with DNA extractions from mouse feces. This work was funded by a grant from National Institutes of Health NIDDK (R01-DK085025 to J.L.S.), an NIH postdoctoral NRSA (5T32AI007328 to A.J.H.), a Stanford University School of Medicine Dean’s Postdoctoral Fellowship (A.J.H.), NSF Graduate Research Fellowships (S.A.S, W.V.T - DGE-114747), an NIH predoctoral NRSA (5T32AI007328 to N.M.D.), and a Smith Stanford Graduate Fellowship (S.A.S.). J.L.S. received an Investigators in the Pathogenesis of Infectious Disease Award from the Burroughs Wellcome Fund.

## Author Contributions

A.J.H., N.M.D., and J.O.G. performed the experiments. D.M.B. conducted blinded scoring, imaging, and analysis of tissue sections. A.J.H., W.V.T., S.A.S., N.M.D., D.M.B., and J.L.S. analyzed and interpreted data, designed experiments, and prepared display items. A.J.H. and J.L.S. wrote the paper. All authors edited the manuscript prior to submission.

